# Machine learning reveals novel targets for both glioblastoma and osteosarcoma

**DOI:** 10.1101/2024.11.05.622056

**Authors:** Nan Li, Max Ward, Muniba Bashir, Yunpeng Cao, Amitava Datta, Zhaoyu Li, Shuang Zhang

## Abstract

Glioblastoma and osteosarcoma originate from the same lineage, yet patients with these two tumour types show significant differences in survival outcomes. Transcriptomic analysis comparing these tumours reveals that over 65% genes show similar expression patterns. Principal component analysis further demonstrates substantial similarities between these two tumour types, albeit with discernible differences. Deep learning analysis employing an autoencoder unveils nuanced distinctions and similarities of these two tumours at a high resolution. A classification model, leveraging gradient boosting with eXtreme Gradient Boosting (XGBoost), achieves high accuracy in distinguishing between these two tumour types. Identification of key contributors to the model’s performance is facilitated by SHapley Additive exPlanations (SHAP), yielding two lists of top target genes with and without considering gender. Notably, these SHAP targets tend to cluster within one or two networks of signalling pathways. Remarkably, gene expression levels of many of these SHAP targets alone can recapitulate survival differences solely based on clinical data between glioblastoma and osteosarcoma patients. Of particular interest, C2ORF72 emerges as a common target from both lists, representing an uncharacterised protein with promising potential as a novel target for diagnostic, prognostic, and therapeutic target for both glioblastoma and osteosarcoma.

## Introduction

The brain, encased within the skull comprising twenty-two bones, is the softest tissue in the human body. Despite its softness, brain tissue has a relatively lower incidence of tumor formation^1^. Glioblastoma, a primary malignant brain tumor, is among the most aggressive tumor types affecting the brain^2^. Surveillance, Epidemiology, and End Results (SEER) data indicate an annual glioblastoma incidence of approximately 3.23 cases per 100,000 individuals, with a higher prevalence among men than women^2^. Glioblastoma has a poor prognosis, with a high mortality rate, and patients typically show poor survival outcomes, with a median survival ranging from 2 to 18% at two years post-diagnosis, and often less than 2.5 years even with intervention^2,3^, underscoring the lack of effective treatment options.

Primary bone tumors, predominantly sarcomas, are rare occurrences, often arising from the soft tissues of bones. Osteosarcoma stands out as the most prevalent primary bone tumor, with an estimated 3,970 diagnoses for the year 2023 in the United States, as per SEER data. Globally, osteosarcoma incidence rates range from 2 to 5 cases per million individuals^4,5^. While patients with osteosarcoma have a 5-year relative survival rate of 60%, this rate drops significantly to 20% for cases that have metastasized to other parts of the body^4,5^.

Given their common origin from ‘soft tissues’ and shared developmental lineage, our objective is to explore the similarities and differences between glioblastoma and osteosarcoma. We intend to achieve this through a comprehensive analysis of genomic data, specifically gene expression profiles obtained from mRNA-Seq data of both tumor types. By leveraging bioinformatics, statistical, deep learning, and machine learning methodologies, we aim to identify novel and common targets between glioblastoma and osteosarcoma, with potential applications in diagnosis, prognosis, and treatment strategies for both cancers.

## Methods

### Genomic data analysis

We obtained mRNA-Seq data for glioblastoma from 153 patients (54 women and 99 men) participating in The Cancer Genome Atlas (TCGA) project, as well as osteosarcoma data from 88 patients (37 women and 51 men) enrolled in the Therapeutically Applicable Research to Generate Effective Treatments (TARGET) project via the GDC portal (NIH, USA). Our dataset exclusively included data from patients with primary tumors. According to the 2021 WHO classifications, the IDH subtypes of these glioblastoma patients from TCGA are 143 IDH-wt cases and 10 IDH-mut cases^6,7^.

To minimize the sequencing bias between TCGA and TARGET datasets, all raw counts were normalized using four housekeeping genes, GAPDH, ACTB, PUM1, and UBC as suggested from pan-cancer studies^8,9^. We performed DESeq2^10^ analysis to compare gene expression counts derived from mRNA-Seq data between glioblastoma and osteosarcoma, as well as within each gender, utilizing default parameters. Transcripts or genes having at least a 2-fold expression difference and adjusted p-values below 0.05 were deemed significantly differentially expressed. Subsequently, heatmaps illustrating selected genes were generated using DESeq2^10^.

### Principle component analysis (PCA)

We performed the PCA using scikit-learn version 1.4.1^11^ on the gene expression data obtained from mRNA-Seq analyses, as described previously, for both glioblastoma and osteosarcoma. Additionally, we collected mRNA-Seq data from 369 hepatocellular carcinoma patients (121 women and 248 men) from TCGA to serve as outliers for comparison.

### Deep learning and machine learning analyses using autoencoder, eXtreme Gradient Boosting (XGBoost), and SHapley Additive exPlanations (SHAP)

We further analyzed the genomic data using deep learning and machine learning algorithms from scikit-learn 1.4.1 with default parameters^11^, incorporating both an autoencoder^12,13^ and XGBoost (version 2.1.2)^14^. The entire datasets were split into two groups: 80% as the training dataset for the modelling and 20% as the validation dataset. The deep autoencoder was built on the Keras 3.0.2 and employed to identify a latent feature space consisting of two features that characterize the data, enabling detection of non-linear relationships. This approach is akin to PCA analysis but is capable of capturing complex, non-linear patterns in the data.

XGBoost^14^, a gradient-boosted trees-based classification model, was utilized to construct random forest tree models. We used the 80-20 training validation split based on a random sample. Subsequently, we employed SHAP (version 0.42.0) to extract Shapley values from the gradient-boosted tree model. This facilitated the identification of genes deemed important to the model’s predictive performance. Top target genes were determined by ranking genes according to their SHAP-computed values or Shapley values^15,16^. These genes are subsequently referred to as ‘SHAP target genes’.

We used an autencoder to provide an alternative non-linear projection of the data to compare to the PCA. The autoencoder was built using the Keras and Tensorflow deep learning libraries. The input was a flattened to a single dimensional vector containing expression levels for each gene (60660 overall). The encoder and decoder had separate weights, but the mirrored architectures: 3 hidden layers with 256, 128, and 64 features, each fully connected with a ReLu activation function. The middle coding layer compressed down to 2 features using a sigmoid activation function. Two encoded features were picked to match the settings for the PCA. The network was trained in batches of 64 for 100 epochs using the Adam optimizer (learning rate set to 0.001) to minimize a reconstruction loss (measured using MSE). Other settings used the Keras default.

We trained another model to do multi-class classification using the data. The XGBoost model was trained to take a vector of expression levels for each gene as input and produce a classification of tumour type as output. An 80-20 training validation split was done after randomly shuffling the data. A gradient boosted tree model was trained using a maximum tree depth of 3. A soft probability for multiclass loss function was used, and 30 rounds of training was done. Other settings used the XGBoost default.

Subsequently, we employed SHAP to extract Shapley values from the gradient-boosted tree model. This facilitated the identification of genes deemed important to the model’s predictive performance. Top target genes were determined by ranking genes according to their SHAP-computed values or Shapley values. These genes are subsequently referred to as ‘SHAP target genes’.

### Pathway analysis

We conducted the pathway analysis for those SHAP target genes with and without considering gender using Ingenuity (Qiagen) with default parameters.

### Survival analysis

We performed two sets of survival analyses using Kaplan-Meier plots^17^. The first analysis using clinical data alone, while the second analysis employed gene expression data exclusively. For the overall survival analysis, clinical data from glioblastoma and osteosarcoma patients sourced from TCGA and TARGET were utilized. The ‘days to death’ parameter was employed to calculate survival time, while ‘days to last follow-up’ was used as censored time for patients who were still alive at the time of analysis.

Furthermore, we conducted an overall survival analysis for both glioblastoma and osteosarcoma patients collectively. This analysis involved segregating all cancer patients based solely on the expression levels of SHAP target genes, categorizing them as either greater or smaller than the median values.

To assess the similarities between clinical data-based survival curves and gene expression-based survival curves, we employed Kullback-Leibler divergence (KL divergence)^18^. KL values below 0.001 were interpreted as indicative of a high degree of similarity between the two curves.

We also conducted the progression-free survival (PFS) analysis for these targets and the PFS data showed similar patterns with the overall survival data (data not shown).

## Results

### Highly similar expression levels for the majority of genes between glioblastoma and osteosarcoma

We collected mRNA-Seq data of glioblastoma from 156 patients in the TCGA project and also osteosarcoma from 88 patients in the TARGET project (Figure 1A). Utilizing DESeq2 analysis^10^, we compared gene expression counts from mRNA-Seq data between glioblastoma and osteosarcoma. Our findings indicated that out of a total of 60,600 transcripts and 19,962 protein-coding genes analyzed, approximately 21,007 transcripts and 6,432 genes showed significant differential expression with at least 2-fold differences (Figure S1A). Remarkably, the expression levels of the remaining 39,653 transcripts and 13,530 genes were found to be similar between glioblastoma and osteosarcoma, constituting approximately 65.4% of total transcripts and 67.8% of total genes, respectively (Figure 1B).

**Figure 1.**
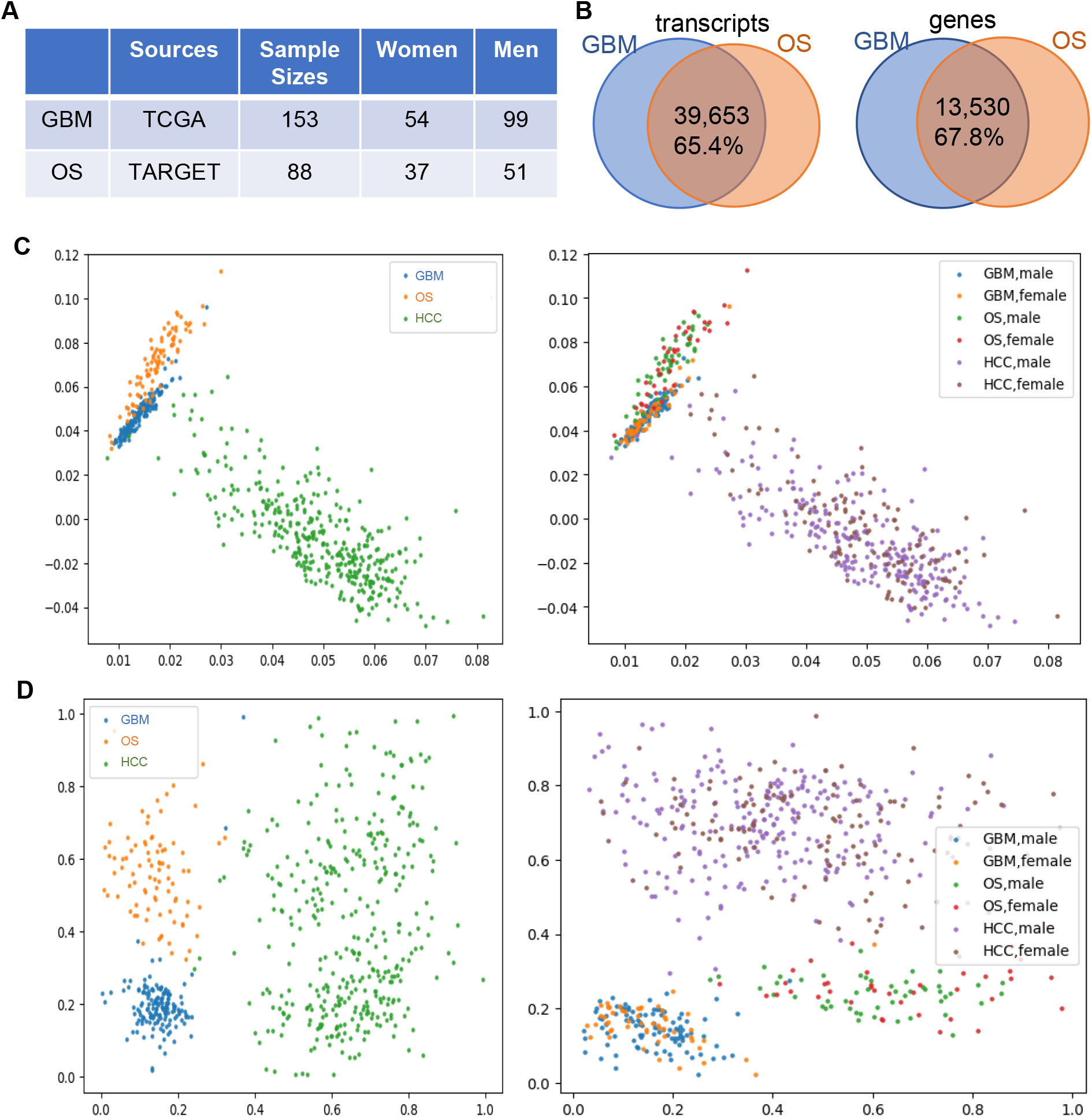
Gene expression patterns between glioblastoma (GBM) and osteosarcoma (OS) patients. (A) Sample sizes and resources. (B) Similarly expressed transcripts and genes between GBM and OS patients using DESeq2 with fold changes < 2 and adjusted P values >= 0.05. (C, D) The principal component analysis (PCA (C) and deep learning autoencoder analysis (D) of gene expression between GBM and OS patients. Hepatocellular carcinoma (HCC) data from TCGA was used as outliers. Left, comparison without considering genders; right, comparison with genders.

Among the 13,530 non-significantly differentially expressed genes, 444 were identified as silenced genes, and 107 genes displayed almost identical expression levels between the two tumor types. Even among the differentially expressed transcripts and 6,432 genes, the majority (5,287 genes) showed higher expression levels above the median in both tumor types, with only 1,045 genes expressing less than the median counts. Heatmap analysis further demonstrated similar expression patterns for these highly expressed 5,287 genes between glioblastoma and osteosarcoma (Figure S1C), suggesting the presence of similarities in differentially expressed genes between the two tumor types. When considering this aspect, the overall similarities in gene expression between glioblastoma and osteosarcoma could potentially exceed 94%.

Additionally, as shown in Figure S1B, a comparison of gene expression data for each gender between glioblastoma and osteosarcoma yielded similar results to those obtained without considering gender (Figure S1A), indicating that the observed similarities in gene expression between the two tumor types are independent of gender.

### Validation of similarities and differences in gene expression between glioblastoma and osteosarcoma patients by PCA, deep learning, and machine learning

To compare gene expression levels across individual patients between glioblastoma and osteosarcoma, we conducted PCA analysis, utilizing gene expression data of hepatocellular carcinoma from TCGA as outliers. As illustrated in Figure 1C, the PCA results revealed strikingly similar expression patterns between glioblastoma and osteosarcoma patients, albeit with discernible differences, which were notably distinct from those observed in hepatocellular carcinoma patients. To further validate these findings, we employed a deep learning analysis, an autoencoder, to extract two latent features for all tumor types. The results (Figure 1D) showcased detailed insights into the similarities and differences between glioblastoma and osteosarcoma. While both tumor types showed similar values along the X axis, Osteosarcoma demonstrated higher values along the Y axis compared to glioblastoma, suggesting their classification under the same tumor category but with notable distinctions. Notably, interpreting the distance across the autoencoder axes proved challenging. However, it is worth noting that while two axes were necessary to differentiate glioblastoma from osteosarcoma, only one was needed to separate both tumor types from hepatocellular carcinoma.

To further elucidate the key drivers underlying the similarities and differences in gene expression between glioblastoma and osteosarcoma, we constructed a classification model using XGBoost^14^ from the gene expression data of each tumor type. We chose XGBoost since it implements gradient boosted trees. This has the advantage of being interpretable. Remarkably, the model showed a high accuracy of 0.98, underscoring its efficacy in distinguishing between the two tumor types. The Shapley value calculation for trees is converges on high confidence estimates rapidly as compared to other models like neural networks. Subsequently, employing SHAP^15,16^ analysis to compute Shapley values for all genes, we identified a set of genes contributing to the observed similarities and differences between glioblastoma and osteosarcoma (Figure 2A).

**Figure 2.**
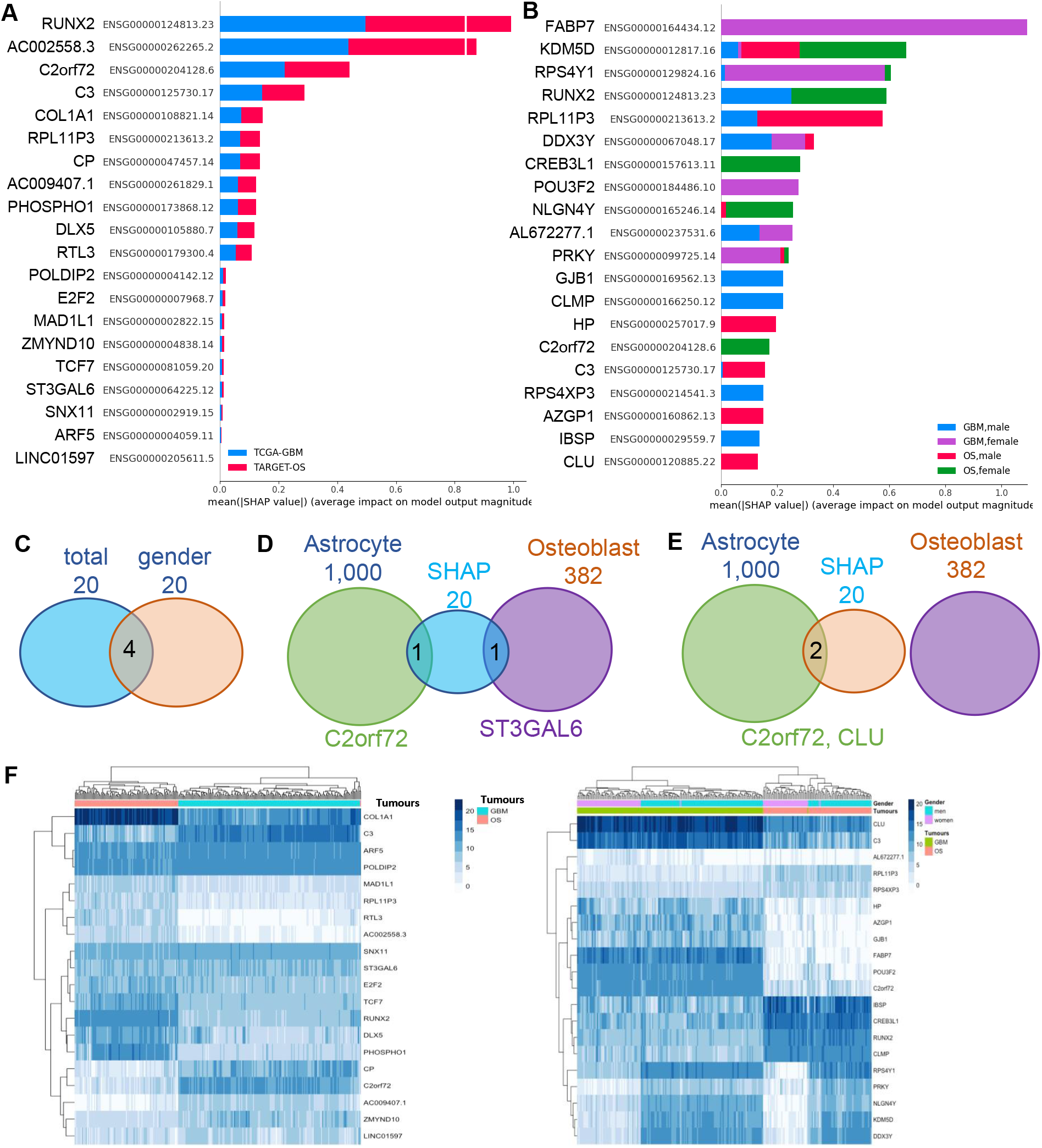
SHAP targets from machine learning. (A, B) Top SHAP targets were identified based on Shapley values obtained from the gradient boosted tree model built using gene expression data between glioblastoma (GBM) and osteosarcoma (OS) patients with (B) and without (A) considering genders. (C) Four common targets between comparisons with and without considering genders. (D, E) Common target genes between signature genes from astrocytes and osteoblast cells and SHAP targets with (E) and without (D) considering genders. (F) Heatmaps of gene expression for SHAP targets with (right) and without (left) considering genders.

We also conducted analogous analyses while considering gender. However, both PCA and autoencoder analyses revealed no discernible differences compared to the results obtained without considering gender (Figures 1C and 1D). Notably, the inclusion of genders in XGBoost models resulted in an accuracy score of 0.99. Despite this high accuracy, SHAP analysis revealed a distinct list of target genes compared to those obtained without considering genders (Figure 2B).

Further analysis revealed four genes present in both lists of SHAP targets, regardless of gender considerations (Figure 2C), including ENSG00000124813.23 (RUNX2), ENSG00000125730.17 (C3), ENSG00000204128.6 (C2ORF72), and ENSG00000213613.2 (RPL11P3). To delve deeper into these targets, we collated 1,000 and 382 signature genes of astrocytes and osteoblast cells^19,20^, respectively, from which glioblastoma and osteosarcoma originate. These signature genes, previously reported from single-cell RNA sequencing studies of human brain and bone marrow^19,20^, overlapped with some SHAP targets, further elucidating their relevance (Figures 2D and 2E). Those SHAP targets without considering genders were overlapped with one of signature genes in astrocytes (C2ORF72) and one in osteoblast cells (ST3GAL6), respectively (Figure 2D). For those SHAP targets with considering genders, only two of them were overlapped with signature genes in astrocytes (C2ORF72 and CLU) but none with osteoblast cells (Figure 2E).

Finally, heatmap visualization of the expression of both lists of SHAP target genes revealed distinct patterns. Some genes were specific to individual tumor types (e.g., COL1A1 and C3 in the list without considering genders), while others showed similarities between the two tumor types (e.g., POLDIP2, ARF5, and SNX11 in the same list).

Additionally, some genes demonstrated similarities between the tumor types but were specific to each gender (e.g., NLGN4Y, KDM5D, and DDX3Y in the list considering genders) (Figure 2F).

### The pathway analyses of SHAP target genes

We conducted Ingenuity pathway analyses for SHAP target genes, both with and without considering genders. Surprisingly, we observed that 13 out of 20 SHAP targets from the list without considering genders were implicated in a single network of signaling pathways (Figure 3A), while 17 out of 20 SHAP targets from the list with genders were associated with two distinct networks of signaling pathways (Figures 3B and 3C). These SHAP targets were interconnected through classical signaling pathways such as PKA/ERK, AKT, CREB, and/or involved in epigenetic regulation through P300 and histones.

**Figure 3.**
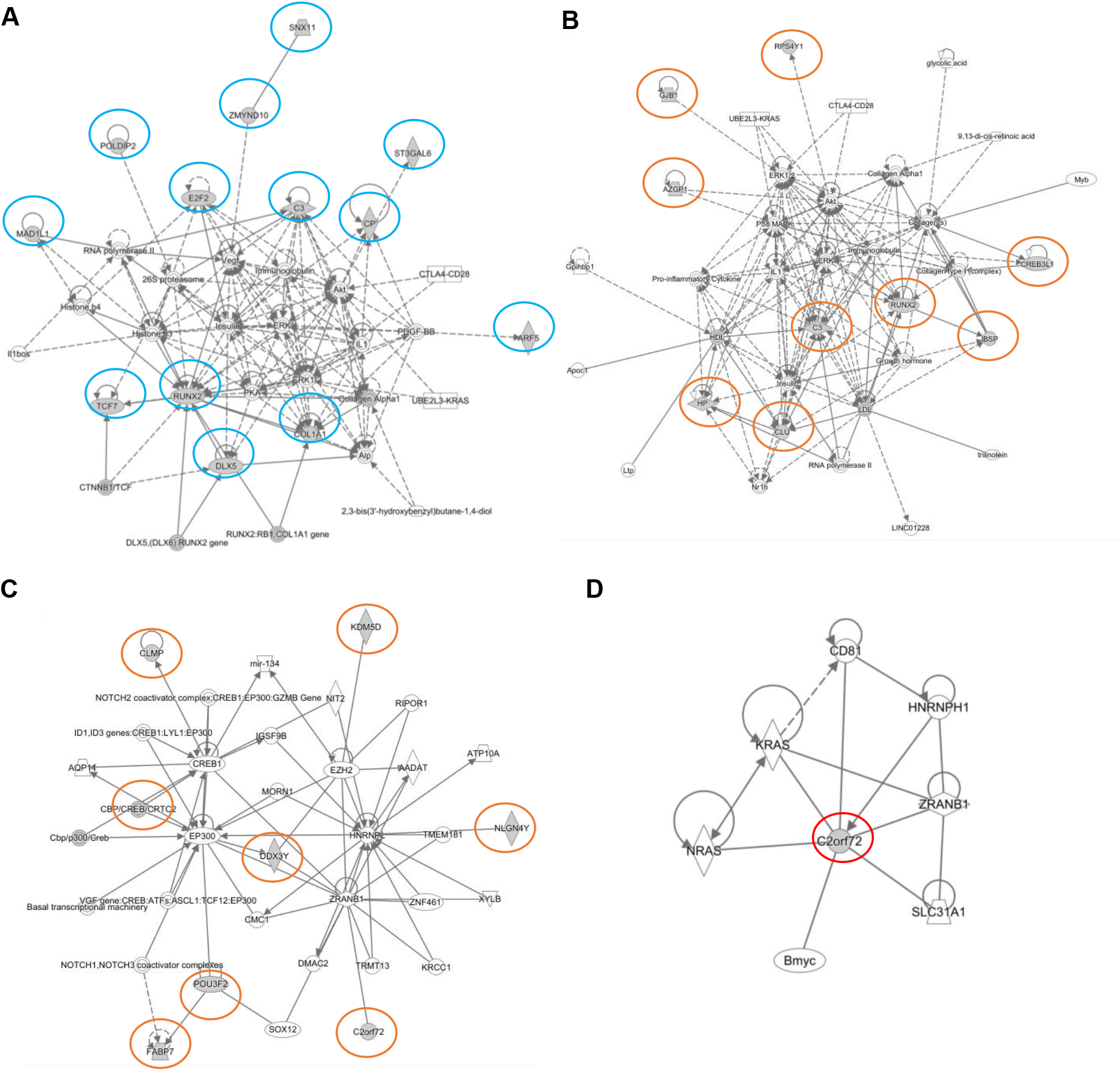
The pathway analysis of SHAP targets using Ingenuity. (A) The pathways for SHAP targets (highlighted in circles) without considering genders. (B, C) The pathways of SHAP targets (highlighted in circles) with genders. (D) The pathways for the common SHAP target, C2orf72, in both with and without considering genders.

Moreover, we identified potential pathways associated with the common target, C2ORF72, found in both lists. These pathways included potential connections with oncogenesis pathways such as RAS or MYC, virus entry pathways involving CD81, RNA stability pathways through heterogeneous nuclear ribonucleoprotein HNRNPH1, and de-ubiquitination pathways via ZRANB1 and SLC31A1 (Figure 3D).

### Survival analysis of SHAP target genes

We further conducted survival analysis using clinical data from glioblastoma and osteosarcoma patients. As shown in Figures 4A and 4B, osteosarcoma patients showed markedly superior survival outcomes compared to glioblastoma patients, consistent with previous findings^3-5^. Subsequently, we evaluated the expression levels of SHAP targets, both with and without considering genders (Figures 4C, 4D, S2 and S3).

**Figure 4.**
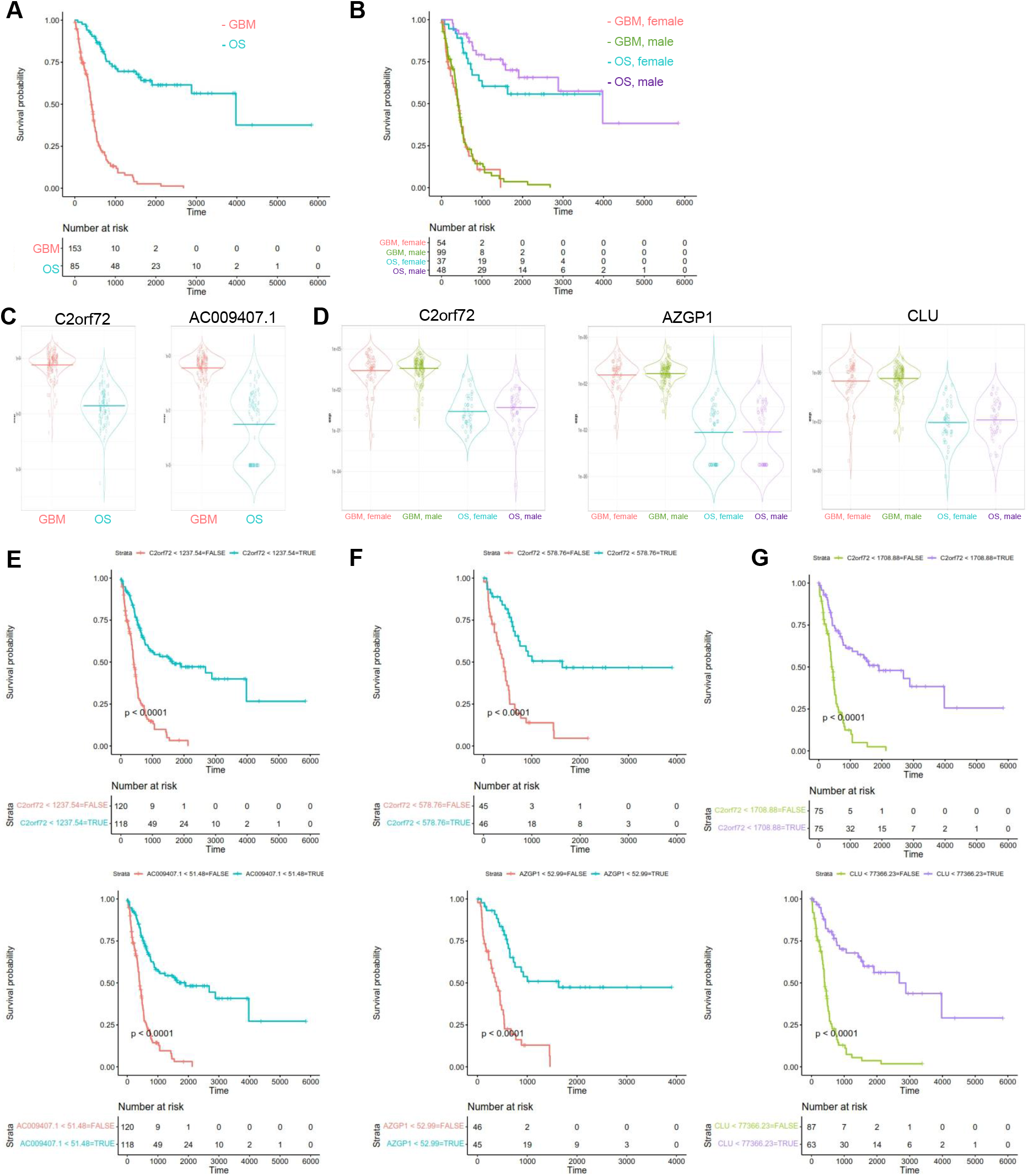
The overall survival analyses. (A, B) The overall survival analysis between glioblastoma (GBM) and osteosarcoma (OS) patients with (B) and without (A) considering genders. (C, D) Expression of selected SHAP target genes between GBM and OS patients with (D) and without (C) considering genders. Y axis, normalised counts. (E-G) The overall survival analysis using the median expression values of selected SHAP target genes among all patients of glioblastoma and osteosarcoma together (E) and in female (F) and male (G) patients.

For instance, in the group without considering genders, the expression levels of C2ORF72 and AC009407.1 were significantly lower in osteosarcoma patients compared to glioblastoma patients (Figure 4C). Notably, C2ORF72 represents an open reading frame of an unknown protein with only one publication regarding the potential regulation of C2ORF72 by microRNA^21^, while AC009407.1 is a long non-coding RNA. Intriguingly, when we conducted survival analysis using the median expression values of C2ORF72 (or AC009407.1) to stratify the entire patient cohort (glioblastoma + osteosarcoma) (Figure 4E), we observed nearly identical survival curves to those generated solely based on clinical data (Figure 4A). By comparing these survival curves with the original ones derived solely from clinical data using KL divergence^18^, we identified a total of eight targets showing similar survival curves, including C2ORF72, AC009407.1, RTL3, LINC01597, CP, COL1A1, C3, and AC009407.1 (Figures 4A and S4). These results underscore the significance of these selected SHAP targets in elucidating the similarities and differences between glioblastoma and osteosarcoma, as observed in PCA and deep learning analyses (Figures 1C and 1D).

Utilizing clinical data, we generated overall survival curves for glioblastoma and osteosarcoma patients based on genders. Remarkably, osteosarcoma patients consistently showed better survival outcomes compared to glioblastoma patients, irrespective of genders, or similar trends observed between males and females within each cancer type (Figure 4B). Subsequently, we separated the entire patient cohort (glioblastoma + osteosarcoma) into female and male groups and generated survival curves based on the median expression values of SHAP target genes while considering genders (Figures 4F and 4G). In the female group, we found that 60% of SHAP targets (12 genes) could recapitulate the original survival curve derived solely from clinical data (Figure 4B), including C2ORF72, AZGP1, C3, CLU, DDX3Y, FABP7, GJB1, HP, KDM5D, NLGN4Y, POU3F2, and RPS4Y1 (Figures 4F and S5).

Similarly, in the male group, nine SHAP target genes could reproduce the clinical data-based survival curve (Figure 4B), including C2ORF72, CLU, AZGP1, C3, FABP7, GJB1, HP, NLGN4Y, and POU3F2 (Figure 4G and S6), all of which were also included in the female group (Figure 4F and S5).

Combining all survival data with (Figure 4F, 4G, S5 and S6) and without (Figure 4E and S4) considering genders, we observed that C2ORF72 and C3 were common targets across these datasets, suggesting their pivotal role in driving genomic similarities and differences between glioblastoma and osteosarcoma. Furthermore, all nine targets, including C2ORF72, CLU, AZGP1, C3, FABP7, GJB1, HP, NLGN4Y, and POU3F2 (Figure 4F, 4G, S5 and S6), contributed to the similarities observed between genders, while DDX3Y, KDM5D, and RPS4Y1 (Figure 4F and S5) were associated with gender-specific differences.

## Discussions

Investigating the similarities among different tumor types holds promise for identifying common novel diagnostic and prognostic biomarkers and therapeutic targets across diverse cancers. Glioblastoma is possibly originated from mainly astrocytes and their progenitor lineage or astrocytic progeny, such as neural stem cells and other progenitor cells, given that astrocytes are derived from heterogeneous populations of progenitor cells in the neuroepithelium^22-27^. Osteosarcoma mainly originates from osteoblasts. Both glioblastoma and osteosarcoma originate from soft tissues, sharing lineage similarities with astrocytes and osteoblasts, respectively, near bone sites. Our study underscores how these locational similarities manifest certain genomic parallels, suggesting that common genomic features may influence cellular behavior and tumor development in similar anatomical niches.

Astrocytes and osteoblast cells, originating from the central nervous system lineage, serve as precursors for glioblastoma and osteosarcoma, respectively. Consequently, the common targets identified in our machine learning analyses may represent key genes driving the observed genomic and phenotypic similarities between these tumor types.

Notably, only a small fraction of SHAP target genes overlapped with gene signatures of astrocytes and osteoblast cells from single-cell sequencing studies, likely due to the bulk sequencing nature of the mRNA-Seq data used in our analysis. Future investigations employing single-cell sequencing of tumor tissues could yield additional targets overlapping with the original gene signatures of these precursor cells. Comparing gene expression profiles of normal brain and bone tissues could further elucidate the developmental lineage and cellular origins contributing to glioblastoma and osteosarcoma pathogenesis, provided that bulk sequencing data for both tissues become available.

Our analysis of survival curves based solely on the expression levels of SHAP targets suggests a novel set of targets with potential therapeutic implications for improving patient survival in both tumor types. However, rigorous functional validations and further investigations of these SHAP targets are imperative for clinical translation.

Pathway analyses revealed a striking observation: the majority of SHAP targets were implicated in one or two networks of signaling pathways. The ability of machine learning algorithms to discern these targets from thousands of genes underscores the need for comprehensive investigation from computational and biological perspectives. Particularly intriguing are the common SHAP targets, C3 and C2ORF72, which appeared in both lists of SHAP targets, regardless of gender considerations. C3 is a key component of the classical complement system, and studies have demonstrated that complements C3 and C4 are involved in the initiation and progression of tumorigenesis^28^. Elevated C3 expression correlated with poorer survival outcomes for glioblastoma and osteosarcoma patients, suggesting a potential therapeutic strategy of reducing C3 levels to improve patient survival. Conversely, C2ORF72, an uncharacterized protein, holds promise as a novel target for diagnosis, prognosis, and treatment of both tumor types. Future endeavors involving cloning, functional studies, and clinical validations of C2ORF72 are warranted and encouraged.

## Acknowledgements

We thank A/Prof Nathan Pavlos from the School of Biomedical Sciences, University of Western Australia for the helpful discussions and suggestions. We also thank the funding support from the Jack Tiddy Fellowship to Z.L.

## Author Contributions

N.L., M.W., and M.B. conducted the data analysis. Y.C. and A.D. provided guidance on data analysis. Z.L. and S.Z. designed the study.

## Data availability statement

All data used for our study are open access data from TCGA (https://portal.gdc.cancer.gov/analysis_page?app=Downloads).

## Notes

### Competing Interest Statement

The authors have declared no competing interest.

### Summary of Updates

This revision is to only link ORCID to authors

## References

1. Zheng, D., Trynda, J., Williams, C., Vold, J.A., Nguyen, J.H., Harnois, D.M., Bagaria, S.P., McLaughlin, S.A., and Li, Z. (2019). Sexual dimorphism in the incidence of human cancers. BMC Cancer 19, 684. 10.1186/s12885-019-5902-z.

2. Miller, K.D., Ostrom, Q.T., Kruchko, C., Patil, N., Tihan, T., Cioffi, G., Fuchs, H.E., Waite, K.A., Jemal, A., Siegel, R.L., and Barnholtz-Sloan, J.S. (2021). Brain and other central nervous system tumor statistics, 2021. CA Cancer J Clin 71, 381–406. 10.3322/caac.21693.

3. Poon, M.T.C., Sudlow, C.L.M., Figueroa, J.D., and Brennan, P.M. (2020). Longer-term (>/= 2 years) survival in patients with glioblastoma in population-based studies pre- and post-2005: a systematic review and meta-analysis. Sci Rep 10, 11622. 10.1038/s41598-020-68011-4.

4. Sadykova, L.R., Ntekim, A.I., Muyangwa-Semenova, M., Rutland, C.S., Jeyapalan, J.N., Blatt, N., and Rizvanov, A.A. (2020). Epidemiology and Risk Factors of Osteosarcoma. Cancer Invest 38, 259–269. 10.1080/07357907.2020.1768401.

5. Smeland, S., Bielack, S.S., Whelan, J., Bernstein, M., Hogendoorn, P., Krailo, M.D., Gorlick, R., Janeway, K.A., Ingleby, F.C., Anninga, J., et al. (2019). Survival and prognosis with osteosarcoma: outcomes in more than 2000 patients in the EURAMOS-1 (European and American Osteosarcoma Study) cohort. Eur J Cancer 109, 36–50. 10.1016/j.ejca.2018.11.027.

6. Zakharova, G., Efimov, V., Raevskiy, M., Rumiantsev, P., Gudkov, A., Belogurova-Ovchinnikova, O., Sorokin, M., and Buzdin, A. (2022). Reclassification of TCGA Diffuse Glioma Profiles Linked to Transcriptomic, Epigenetic, Genomic and Clinical Data, According to the 2021 WHO CNS Tumor Classification. Int J Mol Sci 24. 10.3390/ijms24010157.

7. Louis, D.N., Perry, A., Wesseling, P., Brat, D.J., Cree, I.A., Figarella-Branger, D., Hawkins, C., Ng, H.K., Pfister, S.M., Reifenberger, G., et al. (2021). The 2021 WHO Classification of Tumors of the Central Nervous System: a summary. Neuro Oncol 23, 1231–1251. 10.1093/neuonc/noab106.

8. Krasnov, G.S., Kudryavtseva, A.V., Snezhkina, A.V., Lakunina, V.A., Beniaminov, A.D., Melnikova, N.V., and Dmitriev, A.A. (2019). Pan-Cancer Analysis of TCGA Data Revealed Promising Reference Genes for qPCR Normalization. Front Genet 10, 97. 10.3389/fgene.2019.00097.

9. Consortium, I.T.P.-C.A.o.W.G. (2020). Pan-cancer analysis of whole genomes. Nature 578, 82–93. 10.1038/s41586-020-1969-6.

10. Love, M.I., Huber, W., and Anders, S. (2014). Moderated estimation of fold change and dispersion for RNA-seq data with DESeq2. Genome Biol 15, 550. 10.1186/s13059-014-0550-8.

11. Fabian, P., Gaël, V., Alexandre, G., Vincent, M., Bertrand, T., Olivier, G., Mathieu, B., Peter, P., Ron, W., Vincent, D., et al. (2011). Scikit-learn: Machine Learning in Python. J. Mach. Learn. Res. 12, 2825–2830.

12. Kramer, T.H. (1991). Application of nonlinear regression to the estimation of in vitro antagonist dissociation constants and relative potency problems. Proc West Pharmacol Soc 34, 425–427.

13. Liou, C., Cheng, W., Liou, J., and Liou, D. (2014). Autoencoder for words. Neurocomputing 139, 84–96.

14. Tianqi, C., and Carlos, G. (2016). XGBoost: A Scalable Tree Boosting System. Proceedings of the 22nd ACM SIGKDD International Conference on Knowledge Discovery and Data Mining. Association for Computing Machinery.

15. Najmi, M.S.a.A. (2020). The many Shapley values for model explanation. arXiv, 1908.08474.

16. Scott, M.L., and Su-In, L. (2017). A unified approach to interpreting model predictions. Proceedings of the 31st International Conference on Neural Information Processing Systems. Curran Associates Inc.

17. Kaplan, E.L., and Meier, P. (1958). Nonparametric Estimation from Incomplete Observations. Journal of the American Statistical Association 53, 457–481. 10.1080/01621459.1958.10501452.

18. Kullback, S., and Leibler, R.A. (1951). On Information and Sufficiency. The Annals of Mathematical Statistics 22, 79–86, 78.

19. McKenzie, A.T., Wang, M., Hauberg, M.E., Fullard, J.F., Kozlenkov, A., Keenan, A., Hurd, Y.L., Dracheva, S., Casaccia, P., Roussos, P., and Zhang, B. (2018). Brain Cell Type Specific Gene Expression and Co-expression Network Architectures. Sci Rep 8, 8868. 10.1038/s41598-018-27293-5.

20. Lee, N.Y.S., Li, M., Ang, K.S., and Chen, J. (2023). Establishing a human bone marrow single cell reference atlas to study ageing and diseases. Front Immunol 14, 1127879. 10.3389/fimmu.2023.1127879.

21. Khorasani, M., Shahbazi, S., Hosseinkhan, N., and Mahdian, R. (2019). Analysis of Differential Expression of microRNAs and Their Target Genes in Prostate Cancer: A Bioinformatics Study on Microarray Gene Expression Data. Int J Mol Cell Med 8, 103–114. 10.22088/IJMCM.BUMS.8.2.103.

22. Liu, C., Sage, J.C., Miller, M.R., Verhaak, R.G., Hippenmeyer, S., Vogel, H., Foreman, O., Bronson, R.T., Nishiyama, A., Luo, L., and Zong, H. (2011). Mosaic analysis with double markers reveals tumor cell of origin in glioma. Cell 146, 209–221. 10.1016/j.cell.2011.06.014.

23. Alcantara Llaguno, S., Chen, J., Kwon, C.H., Jackson, E.L., Li, Y., Burns, D.K., Alvarez-Buylla, A., and Parada, L.F. (2009). Malignant astrocytomas originate from neural stem/progenitor cells in a somatic tumor suppressor mouse model. Cancer Cell 15, 45–56. 10.1016/j.ccr.2008.12.006.

24. Alcantara Llaguno, S., Sun, D., Pedraza, A.M., Vera, E., Wang, Z., Burns, D.K., and Parada, L.F. (2019). Cell-of-origin susceptibility to glioblastoma formation declines with neural lineage restriction. Nat Neurosci 22, 545–555. 10.1038/s41593-018-0333-8.

25. Wang, Z., Sun, D., Chen, Y.J., Xie, X., Shi, Y., Tabar, V., Brennan, C.W., Bale, T.A., Jayewickreme, C.D., Laks, D.R., et al. (2020). Cell Lineage-Based Stratification for Glioblastoma. Cancer Cell 38, 366–379e368. 10.1016/j.ccell.2020.06.003.

26. Hu, Y., Jiang, Y., Behnan, J., Ribeiro, M.M., Kalantzi, C., Zhang, M.D., Lou, D., Haring, M., Sharma, N., Okawa, S., et al. (2022). Neural network learning defines glioblastoma features to be of neural crest perivascular or radial glia lineages. Sci Adv 8, eabm6340. 10.1126/sciadv.abm6340.

27. Ah-Pine, F., Khettab, M., Bedoui, Y., Slama, Y., Daniel, M., Doray, B., and Gasque, P. (2023). On the origin and development of glioblastoma: multifaceted role of perivascular mesenchymal stromal cells. Acta Neuropathol Commun 11, 104. 10.1186/s40478-023-01605-x.

28. Thurman, J.M., Laskowski, J., and Nemenoff, R.A. (2020). Complement and Cancer-A Dysfunctional Relationship? Antibodies (Basel) 9. 10.3390/antib9040061.

