## supplementary figures for "Machine learning reveals novel targets for both glioblastoma and osteosarcoma"

**A**

| | Transcripts | Genes |
|---|---|---|
| Total | 60,660 | 19,962 |
| SDE | 21,007 | 6,432 |
| > median | 11,435 | 5,287 |
| < median | 9,572 | 1,045 |
| Non-SDE | 39,653 | 13,530 |
| silenced | 2,731 | 444 |
| identical | 190 | 107 |

**B**

| | Women | | Men | |
|---|---|---|---|---|
| | Transcripts | Genes | Transcripts | Genes |
| Total | 60,660 | 19,962 | 60,660 | 19,962 |
| SDE | 22,216 | 7,204 | 21,733 | 7,326 |
| > median | 11,109 | 5,994 | 10,868 | 6,175 |
| < median | 11,107 | 1,210 | 10,865 | 1,151 |
| Non-SDE | 38,444 | 12,758 | 38,927 | 12,636 |
| silenced | 3,274 | 486 | 3,156 | 470 |
| identical | 267 | 118 | 240 | 114 |

**C**

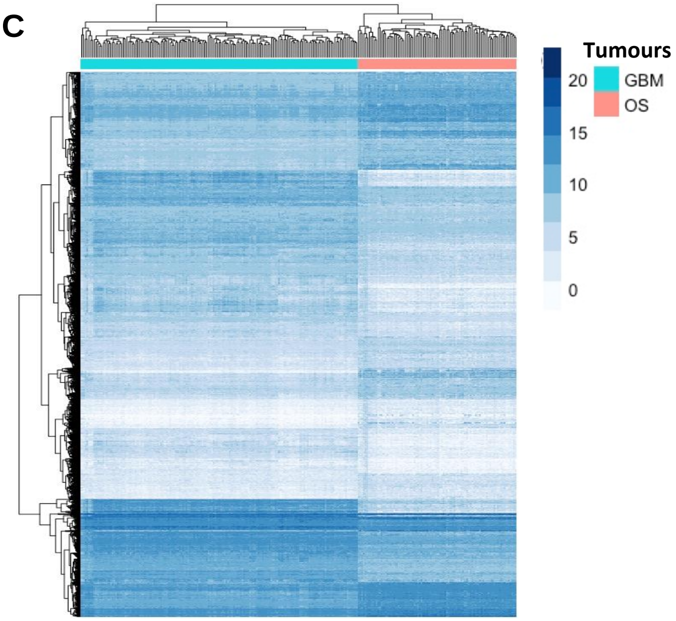


**Figure S1 Similarities in gene expression between glioblastoma (GBM) from TCGA and osteosarcoma (OS) and from TARGET.**

(A) Comparison of gene expression between GBM and OS patients using DESeq2. SDE, significantly differentially expressed genes with fold changes >= 2 and adjusted P values < 0.05. Identical, equally expressed genes with fold changes > -1.005 and also < 1.005.

(B) Comparison of gene expression between GBM and OS patients in each gender.

(C) Heatmaps of gene expression for those highly and differentially expressed genes between GBM) and OS patients without considering genders and also greater than the median values.

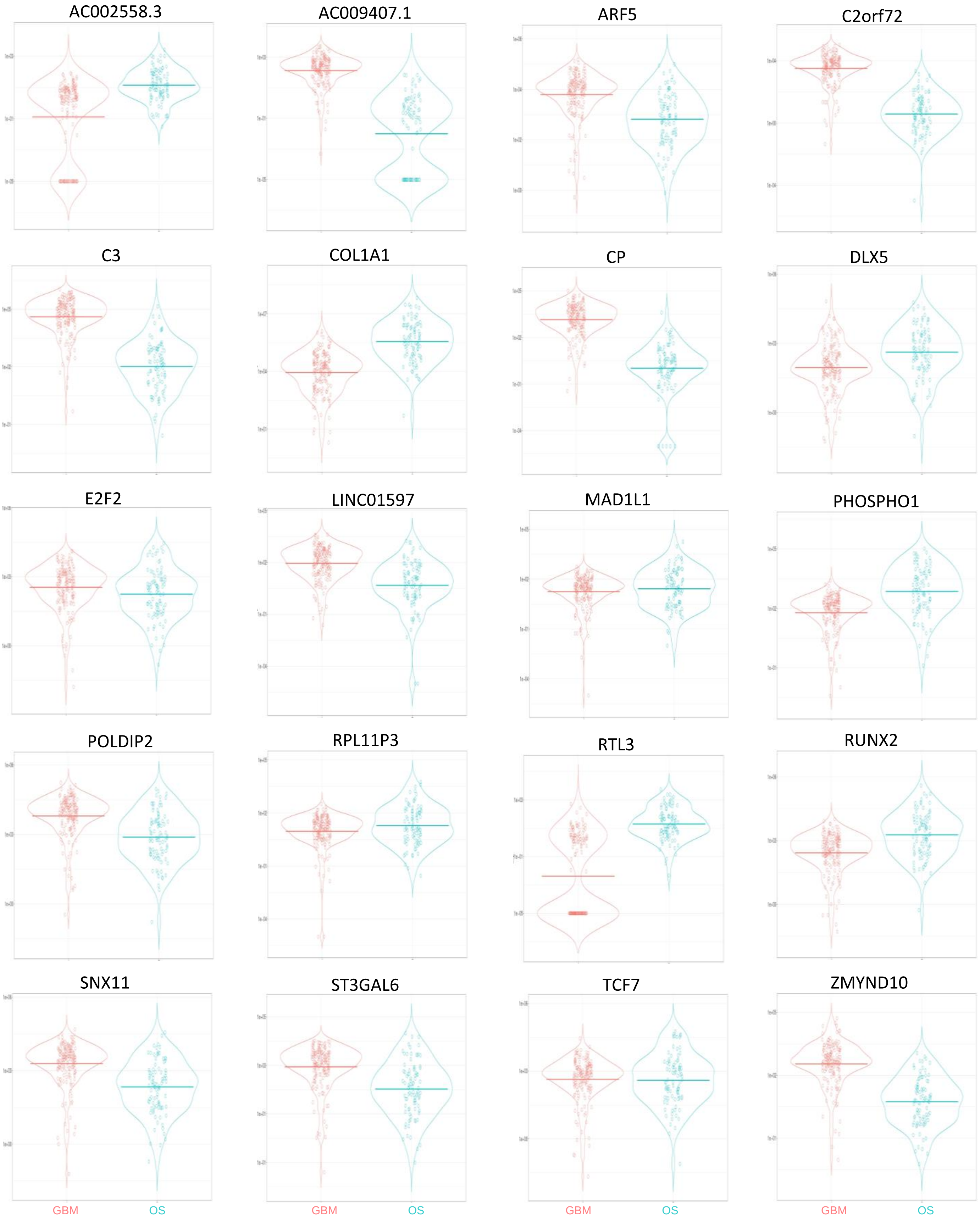


**Figure S2 The expression levels of SHAP genes between glioblastoma (GBM) and osteosarcoma (OS) patients without considering genders.**
Y axis, normalised counts.

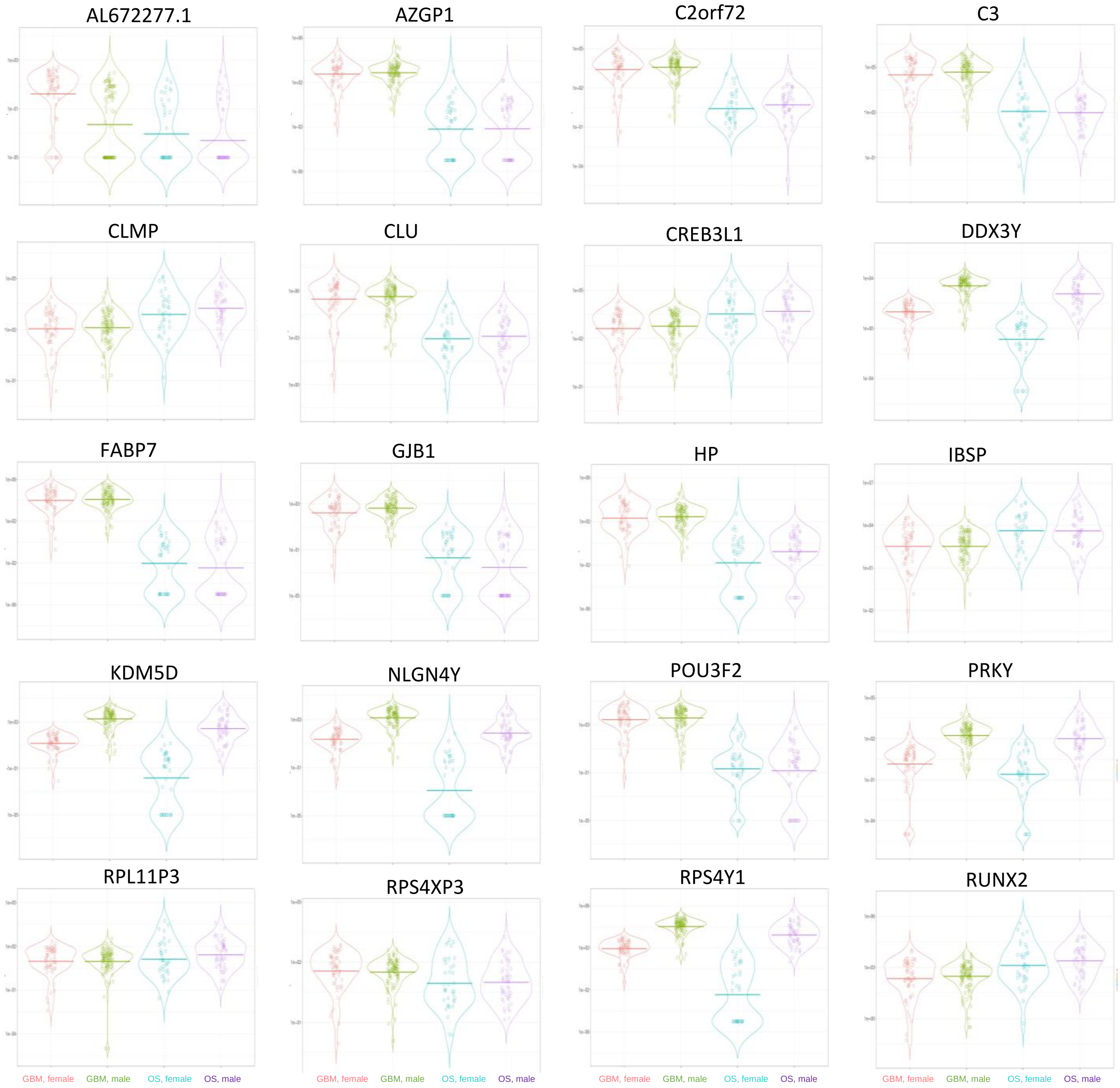


**Figure S3 The expression levels of SHAP genes between glioblastoma (GBM) and osteosarcoma (OS) patients with genders.**
Y axis, normalised counts.

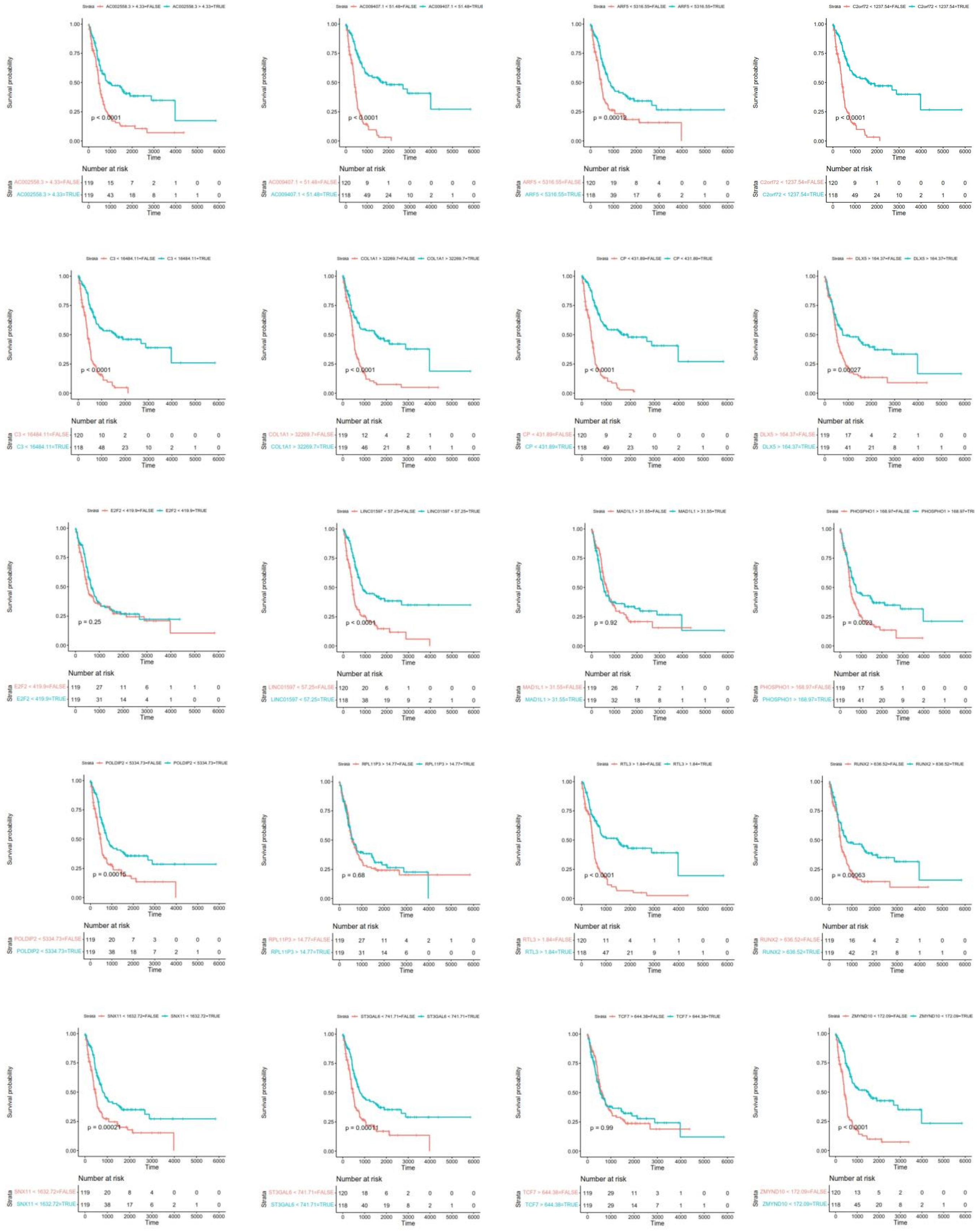

**Figure S4 The overall survival analysis between glioblastoma (GBM) and osteosarcoma (OS) patients without considering genders.**

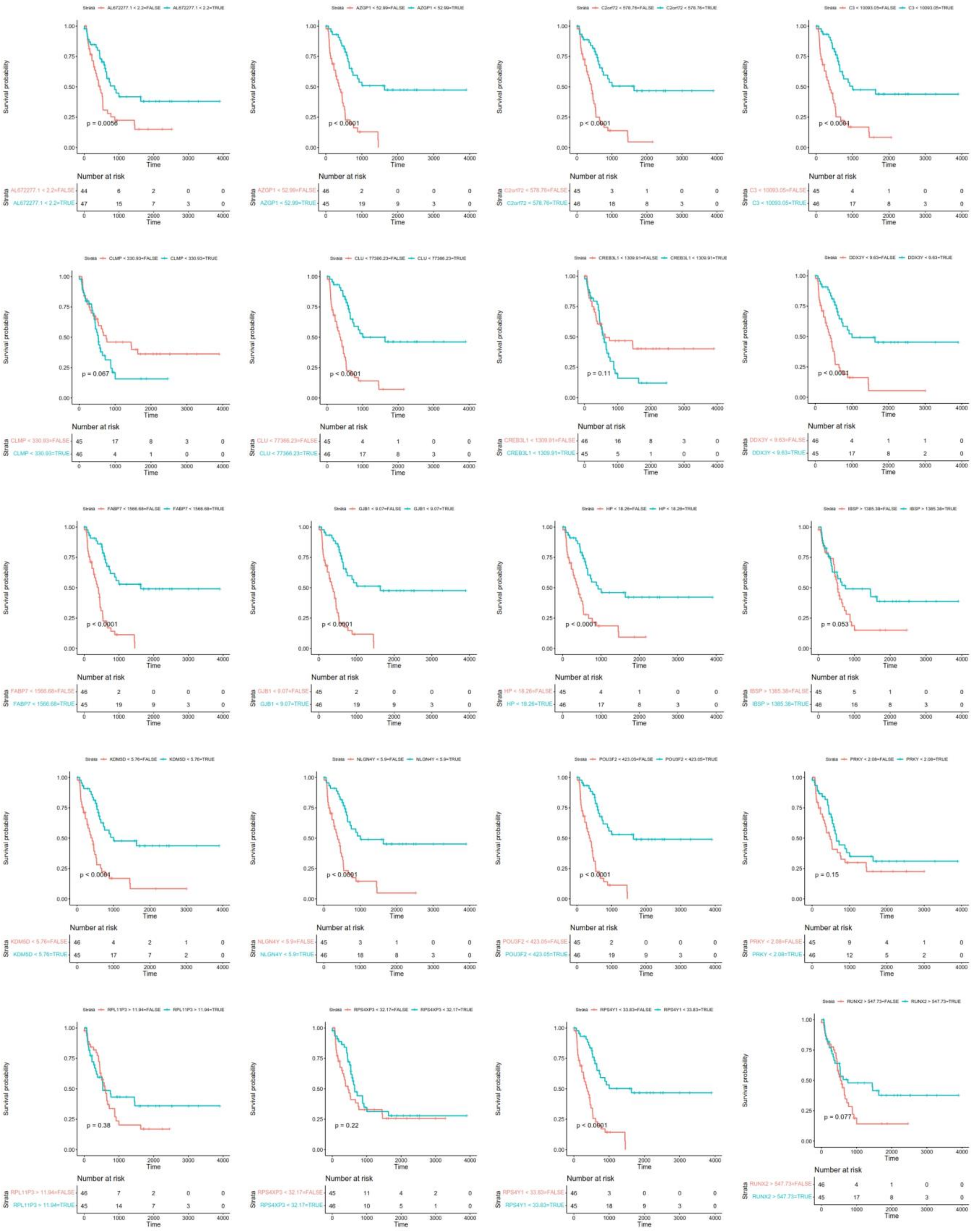

**Figure S5 The overall survival analysis between female glioblastoma (GBM) and osteosarcoma (OS) patients.**

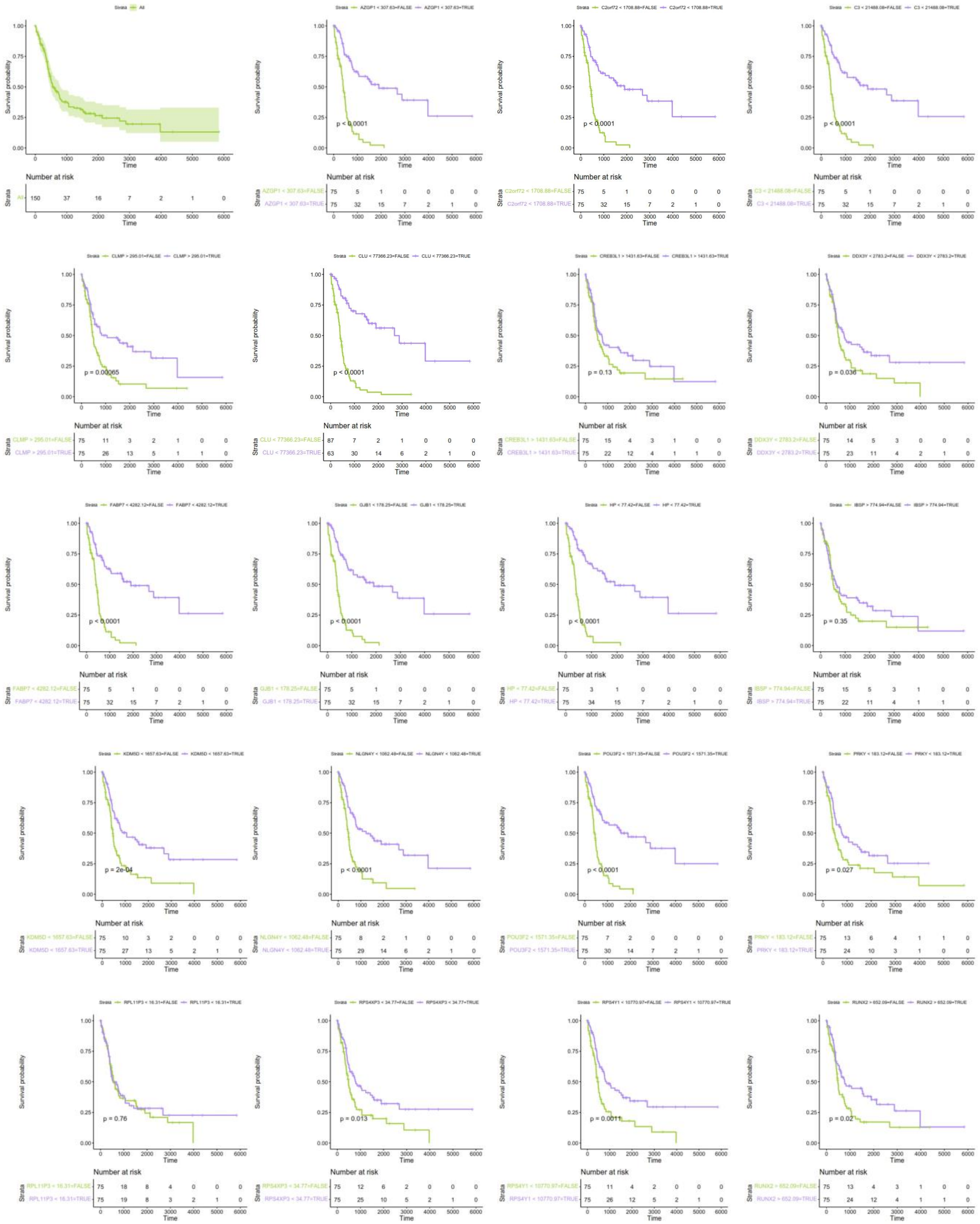


**Figure S6 The overall survival analysis between male glioblastoma (GBM) and osteosarcoma (OS) patients.**
